# EXCON: a scalable, reproducible Nextflow pipeline for CAFE-based gene EXpansion and CONtraction analysis

**DOI:** 10.64898/2026.09.25.754512

**Authors:** Christopher Wyatt, Fernando Duarte, Alexandra Cerqueira De Araujo, Simon Murray, Cintia Oi, Seirian Sumner

## Abstract

Gene family expansion and contraction analysis is a commonly used approach in evolutionary comparative genomics to identify lineage-specific evolutionary pressures on gene copy/family number. CAFE is one of the most popular tools to measure changes in gene family size, but requires a high level of bioinformatics expertise, manual parameter tuning and preprocessing. Here we present EXCON, an automated, Eco-Flow community-developed Nextflow pipeline that integrates NCBI genome download, annotation quality and format pre-processing, orthologous gene family identification (Orthofinder), ultrametric tree formatting, birth-death modelling (CAFE), and GO enrichment analysis (eggNOG-mapper/TopGO) into a single, portable workflow. EXCON accepts genome assemblies and annotations, supports local, HPC and cloud-based computing environments, and requires minimal user configuration. The pipeline auto-selects appropriate CAFE lambda and error models, and separates high variance orthogroups for separate treatment. Applied to a 24-species primate dataset spanning Catarrhini, Platyrrhini and Strepsirrhini, the pipeline identified 358 rapidly evolving gene families, including a DUX (double homeobox) gene showing repeated, independent copy-number turnover across multiple lineages, consistent with the known instability of the D4Z4 macrosatellite repeat it resides in. EXCON is freely available at https://github.com/Eco-Flow/excon.

## Introduction

The number of copies of a gene in a genome is not fixed across species. Gene families expand through duplication and contract through loss, and these changes in copy number are a primary mechanism by which lineages adapt to novel ecological niches, evolve complex phenotypes(1, 2), and respond to arms-race dynamics between hosts and pathogens(3, 4). Identifying which gene families have expanded or contracted along specific branches of a phylogeny and quantifying the statistical confidence of those changes is therefore a fundamental goal of comparative genomics.

The Computational Analysis of gene Family Evolution (CAFE) framework (5) models gene family size changes as a stochastic birth-death process along a phylogenetic tree, providing maximum-likelihood estimates of the rate of gene gain and loss (the lambda parameter: λ) and identifying gene families that evolve significantly faster or slower than the genome-wide average(6, 7). Since its introduction, CAFE has been applied across the tree of life, including mammals(8, 9), insects(10, 11) and plants(12, 13); and version 5 of the software (CAFE5) introduced gamma rate categories to model among-family rate variation(5).

Despite the maturity of the underlying statistical framework, performing a complete CAFE analysis from raw genome data remains technically demanding. A typical analysis requires: downloading and standardising genome assemblies and annotations from disparate sources; extracting longest protein isoforms; identifying orthologous gene families across all species; constructing and time-calibrating an ultrametric species tree; formatting input files to CAFE’s specific requirements; running multiple model configurations; comparing model likelihoods; and interpreting results in the context of gene function potentially via GO enrichment. Each of these steps involves different tools, file formats, and potential failure debugging, and the lack of an integrated pipeline means that analyses are rarely reproducible across computing environments or research groups(14). Currently, no tool integrates the complete workflow from genome download through biological interpretation in a portable, community-oriented manner.

Nextflow(15) has emerged as the leading workflow management system for reproducible bioinformatics, providing native support for containerisation (Docker, Singularity/Apptainer), scalable execution across local machines, HPC clusters, and cloud platforms, and automatic resumption of failed analyses. Further, the worldwide Nextflow community - nf-core(16, 17) - has established a set of best-practice guidelines for Nextflow pipeline development, promoting code quality, testing, and interoperability. These features mean that Nextflow pipelines are adaptable; being modular, they are easily maintained and updated as new versions of different programmes become available. Here we present EXCON (https://github.com/Eco-Flow/excon), a fully automated Nextflow pipeline implementing the complete EXCON analysis workflow following nf-core guidelines. The current release is Version 2.4.0. EXCON addresses key limitations of existing approaches by: (i) automating genome and annotation acquisition from NCBI or user-provided files; (ii) incorporating model convergence detection with progressive differential filtering to handle challenging datasets; (iii) performing automatic gamma/poison model selection across multiple rate categories; (iv) integrating GO enrichment analysis for expanded and contracted gene families; and (v) producing standardised, publication-ready outputs. EXCON is designed for accessibility, requiring only a species list as input, and for reproducibility, with all software dependencies managed through containers.

## Materials and Methods

### Pipeline overview

EXCON is implemented in Nextflow DSL2 and follows nf-core community guidelines (13). The complete workflow comprises major stages including: (i) genome acquisition/download, (ii) quality control (‘STATS’), (iii) pre-processing (gunzip/agat/gffread); (iv) gene ontology (GO) annotation (eggNOG-mapper(18)); (v) orthologous gene family identification (OrthoFinder(19, 20)); (vi) species tree construction and ultrametrification; (vii) birth-death modelling (with CAFE(5)); and (viii) functional enrichment analysis (with TopGO(21)). The pipeline is made up of 40+ modular processes, each containerised independently to ensure software version reproducibility (many of the modules were built by the nf-core community). The input is flexible, allowing the user to bypass the upstream steps entirely by supplying pre-computed OrthoFinder outputs (‘--input_orthogroups’, ‘--input_treè) or pre-computed gene-go hashes (‘--predownloaded_gofiles’).

### Genome acquisition

The pipeline accepts either a two-column CSV file of species names and NCBI RefSeq(22) accession identifiers, and/or a three-column CSV file with paths to local genome FASTA and GFF (Gene Feature Format) annotation files. Examples of input sample sheets can be found with the repository (https://github.com/Eco-Flow/excon/blob/main/data/input_*.csv). When NCBI accession identifiers are provided, genomes and annotations are downloaded using NCBIGenomeDownload (v0.3.3)(23). A mixture of both input line types is also accepted by the pipeline.

### Quality control

Optional quality control statistics are computed using BUSCO v6.0.0(24), QUAST v5.3.0(25), and AGAT v1.6.1(26) (‘agat_sp_statistics.pl’) when the ‘--stats’ parameter is specified, providing assembly completeness, contiguity, and annotation summary metrics, which can be used to determine if the genomes are of sufficient quality to be comparable. We also used the pipeline nf-core/genomeqc(27) to ensure that our sets of genomes looked comparable. For this we chose genomes with ≥95% complete BUSCOs, with no obvious evidence of excessive duplication or fragmented annotation, but previous papers have used other thresholds between 70%(28) to 95%(29).

### Preprocessing

GFF/GTF files were filtered to one longest-coding isoform per gene with AGAT v1.6.1 (‘agat_sp_keep_longest_isoform.pl’)(26). Protein sequences were extracted using GFFRead v0.12.7(30) (‘-y -S’). A custom RENAME_FASTA module/script mapped transcript to gene IDs across NCBI, Ensembl and BRAKER/AUGUSTUS naming conventions, giving GFFread and AGAT outputs matching gene-level identifiers, and resolved premature in-frame stop codons via one of three modes (‘-- internal_stop_action’): strip (splice out all stops), drop (discard the gene) or longest_orf (default; longest ORF between stops). Affected genes were logged per species.

### Orthologous gene family identification

Orthologous gene families are identified with OrthoFinder v3.1.4(19, 20) or v2.5.5 (‘-- orthofinder_v2’) as an alternative, run on the cleaned proteomes. By default, OrthoFinder performs an all-versus-all DIAMOND v2.1.24.178(31) search, clusters hits into orthogroups by Markov clustering (MCL v22-282(32)) and infers a gene tree per-orthogroup with DendroBLAST, a distance-based method built directly from sequence-similarity scores, without aligning them. Setting ‘--orthofinder_method msà instead builds full alignment-based trees: MAFFT (v7.526(33)) (default) or MUSCLE(34) (‘--orthofinder_msa_prog’), followed by FastTree (v2.2.0(35)) (default), RAxML(36), RAxML-NG or IQ-TREE (with flag ‘--orthofinder_treè). The search, alignment and tree programs can each be set independently via ‘-- orthofinder_search’, ‘--orthofinder_msa_prog’ and ‘--orthofinder_treè. A species tree is inferred from the gene trees with STAG and rooted with STRIDE (Species_Tree/SpeciesTree_rooted_node_labels.txt). This tree and its matching gene-count table: Orthogroups/Orthogroups.tsv for v3, or Phylogenetic_Hierarchical_Orthogroups/N0.tsv for v2.5.5 (folded into Orthogroups.tsv from v3.1.0 onward) are passed to CAFE5. All other options are documented in the pipeline README (https://github.com/Eco-Flow/excon).

### Species tree preparation

The input species tree (from OrthoFinder or IQ-TREE) has branch lengths in substitutions per site and is generally non-ultrametric. By default, the module RESCALE_TREE (custom rescale_tree.py) multiplies all branch lengths by ‘-- tree_scale_factor’ (default 1000) to avoid CAFE5 numerical-precision stalls; CAFE_PREP then confirms the tree is rooted and binary (ape(37)) and, only if it remains non-ultrametric, applies chronoMPL(). No rescaling is reapplied, since chronoMPL is linear and doing so would square the initial factor. Where fossil or published node ages are available, ‘--tree_calibrations’ (a TSV of clade, MRCA-defining tips, and minimum/maximum age in Ma) instead directs DATE_TREE to time-calibrate the tree with ape::chronos (configurable model, rate-smoothing λ and rate categories), yielding branch lengths in Myr and bypassing rescaling entirely; convergence warnings and calibration mismatches abort the run. ‘-- input_tree_is_dated’ instead passes an already time-calibrated, ultrametric tree through unchanged. Optionally, ‘--cafe_cladè or ‘--cafe_species’ then restrict the analysis to one clade (named by the MRCA of two or more tips) or an explicit species subset, pruning the tree after any time-calibration, so calibrated node ages are preserved before it reaches CAFE_PREP. In all cases, tree tips and gene-count columns have any .clean suffix (from Preprocessing) stripped, and any remaining name mismatch aborts the run.

### CAFE5 model selection

The core birth–death modelling stage runs as a series of CAFE5 calls tied together by four modules. Before any model is fit, HOGs are excluded if they are empty after subsetting to the tree’s species, present in only one species, or reach ≥100 copies in any single species recorded in hog_filtering_report.tsv on every run. CAFE_PREP first fits a single-rate (“Base”) model (‘cafe5 -i hog_gene_counts.tsv -t cafe_input_tree.txt’) to check convergence; since CAFE5 can exit 0 without converging, its log is checked explicitly for a size-differential error, an infinite likelihood, or a missing final-likelihood line, and any of these triggers up to three retries with --cafe_max_differential (default 50, halved each retry) excluding the most variable families. A separate CAFE5 -e call then estimates an error model quantifying assembly/annotation error in the gene counts; if this fails to converge, an empty placeholder is used instead and downstream runs proceed without error correction. Second, CAFE_RUN_K fans out ‘cafe5 -k K -eBase_error_model.txt’ (-k omitted for k=1) in parallel across k = 1–params.cafe_max_k (default 6), the number of discrete gamma rate categories. Third, CAFE_SELECT_K parses each run’s −lnL and selects the k minimising AIC = 2 × n_params + 2 × (−lnL), with n_params = 1 for k = 1 (λ only) and 2 for k > 1 (λ + α); a table of k, −lnL, AIC, BIC and the selected model is written to the results directory (BIC is reported but not used for selection).

Fourth, CAFE_RUN_BEST re-runs only the winning k under a Poisson root-frequency prior (cafe5 -k K -p). This prior is CAFE5’s assumed distribution of ancestral family sizes at the root of the tree: uniform across the observed range by default, or concentrated on smaller, more plausible sizes under the Poisson alternative. The uniform-prior result itself is reused directly from the earlier sweep rather than recomputed. CAFE_MODEL_COMPARE then compares the two −lnL values directly rather than with a significance test, since both models share the same k and parameter count and the lower −lnL is simply the better fit; if only one of the two converged, that one is used automatically, and the process fails only if neither did.

### Convergence detection and progressive filtering

CAFE5 exits with status 0 even when every likelihood evaluation returns infinity, so CAFE_PREP’s base-model run checks its own log for three signatures of non-convergence: a “largest size differential” error; Score (-lnL): inf without a Final -lnL: line; or the Final -lnL: line missing outright. Any of these causes the process to exit 1, triggering a Nextflow retry; up to three retries (hardcoded). From the first retry onward, families are excluded if their max-minus-min copy number across species exceeds a threshold that halves each attempt: --cafe_max_differential (default 50) / 2^(attempt−2), i.e. 50, then 25, then 12. A hog_filtering_report.tsv records every family’s exclusion reason on every attempt. Families excluded specifically for exceeding the differential threshold are written to hog_gene_counts_large.tsv and re-analysed individually by CAFE_RUN_LARGE, each fitting its own independently-estimated λ rather than a shared one, a single shared λ was found not to converge, since no one rate can explain both a modest and an extreme copy-number differential at once (hahnlab/CAFE5#132). The k-sweep (CAFE_RUN_K) has no equivalent recovery: it does not check its own log for non-convergence, so a k value that fails simply produces an unparseable likelihood line and is silently dropped from AIC comparison by CAFE_SELECT_K rather than retried.

MERGE_CAFE_LARGE_RESULTS then stitches every converged per-family result from CAFE_RUN_LARGE into one combined directory and records a per-family λ summary; this step is best-effort (errorStrategy ’ignore’) and its failure does not affect the main CAFE5 run.

### Significant gene family expansion and contraction

CAFE_SIG_FAMILIES reads the best-fit CAFE5 model’s results together with the orthogroup gene-count table and reports, for every node in the tree (tips and internal), each family’s count change and whether that change is significant, using a CAFE5 branch p-value cutoff set by ‘--go_cutoff’ (default 0.05). This is a separate significance call from the one used later for GO enrichment, which applies its own fixed 0.05 threshold. For any node named in ‘--cafe_focus_clades’, the alignment and gene tree behind every significantly expanded family at that node are archived (focus_families.tar.gz) for closer inspection. COMBINE_CAFE_SIG_FAMILIES then merges this with the equivalent results from the large-family track (CAFE_RUN_LARGE), producing one combined table covering every orthogroup CAFE5 was able to fit, whether under the shared k/λ model or individually.

### GO enrichment analysis

Gene-to-GO mappings can either be supplied directly (‘--predownloaded_gofiles’) or generated by eggNOG-mapper within the pipeline (‘--run_eggnog’), which downloads its reference database automatically unless ‘--eggnog_data_dir’ points to a copy already on disk. The annotation step itself can be tuned through the usual alignment thresholds (e-value, bitscore, percent identity, coverage), and restricted to a taxonomic scope if needed. EGGNOG_TO_GO maps these annotations onto gene-level GO terms per species (it looks for mRNA features in the GFF), and EGGNOG_TO_OG_GO rolls that up to the orthogroup level.

CAFE_GO_PREP then pulls out, for every species and internal node, the orthogroups that expanded or contracted with a CAFE5 branch p-value of 0.05 or below. This threshold is fixed, independent of ‘--go_cutoff’. Each species gets its own background set for testing: every orthogroup with at least one gene in that species, not the whole dataset. Sets smaller than 10 orthogroups are dropped rather than tested. CAFE_GO_RUN runs each remaining target against its background through ‘ChopGO_VTS2.pl’, using topGO’s Fisher’s exact test (‘--go_algò, default classic_fisher; the other algorithm options need per-gene scores rather than a membership list, so they return nothing here). No correction for multiple testing is applied unless asked for. ‘--go_typè can switch one on, either family-wise (bonferroni, holm, hochberg, hommel) or FDR (BH, fdr, BY). Results are summarised by functional category and plotted.

’--chromo_gò runs this same machinery again on a different axis: per chromosome or scaffold instead of per significant gene set, to look for functional bias tied to genomic location rather than to expansion or contraction.

### Implementation and availability

EXCON is implemented in Nextflow (v25.10.0 or later) and all software dependencies are containerised using Docker or Singularity/Apptainer, ensuring full reproducibility across computing environments. The pipeline has been tested on local machines, the UCL Myriad HPC cluster (SGE scheduler), local (‘amd’ and ‘arm’ infrastructure) and AWS Batch. Configuration profiles are provided for these environments. Custom institutional configurations can be supplied via ‘-- custom_config’. A report is made at the end of each run (found in results/pipeline_info), that informs the users of cpu, memory and time used per process, as well as other metadata in line with existing nf-core pipelines.

The pipeline follows nf-core(16) guidelines for module structure, parameter schema validation (nf-schema v2.5.1), software version tracking via topic channels, and stub testing. All modules publish outputs to a structured results directory. The pipeline is released under the MIT licence and is available at https://github.com/Eco-Flow/excon.

## Results

### Application to Primate comparative genomics

To demonstrate EXCON’s capabilities on a large, phylogenetically diverse dataset, we applied it to 24 primate species spanning all three major primate radiations: Catarrhini (apes and Old World monkeys, including all extant great ape genera, two gibbon species, four macaque species, baboon, gelada, green monkey, and three colobines), Platyrrhini (marmoset and squirrel monkey), and Strepsirrhini (three lemuriform species and one lorisid). This dataset spans approximately 71 million years of primate evolution, from the Strepsirrhini–Haplorrhini crown divergence to the present(38, 39) and encompasses species with highly variable genome sizes, social organisation, dietary ecology, and life history strategies.

EXCON was run on this dataset using default pipeline settings (nextflow run main.nf -profile test_primate). During preprocessing, a small number of gene models per species contained a premature in-frame stop codon, resolved via the pipeline’s default longest_orf strategy rather than discarding the affected genes (Supplementary Table S1; default is longest_orf, but this is optional using the flag -- internal_stop_action, where a user can drop [remove] these genes if preferred).

OrthoFinder then inferred orthologous gene families and a rooted species tree, and CAFE5 ran successfully with no need for filtering of high variance orthogroups, converging on a birth-death rate of λ = 0.009635 (epsilon: 0.010998, Alpha: 0.7373). Selection of the optimal number of discrete gamma rate categories (k) is a critical but often neglected step in CAFE5 analysis, and here the optimal model was k=2, indicating moderate among-family rate variation: a two-category gamma model captures meaningfully different evolutionary dynamics across gene families, but the absence of support for higher k values (AIC increases monotonically from k=2 onward; **Table 1A**) suggests that rate heterogeneity in this dataset does not require a more complex prior. This is consistent with the relatively close phylogenetic depth of the primate radiation compared to deeper-time comparative datasets. The AIC improvement from k=1 to k=2 was substantial (ΔAIC = 17,848), confirming that a single-rate model would meaningfully underfit these data. Both AIC and BIC are reported in the model selection table, allowing users to apply alternative criteria. At k=2, the Poisson root-frequency model fits the data better than the uniform-prior model (**Table 1B**), with a final “Model Gamma Final Likelihood” (-lnL) of 176160.

**Table 1.**
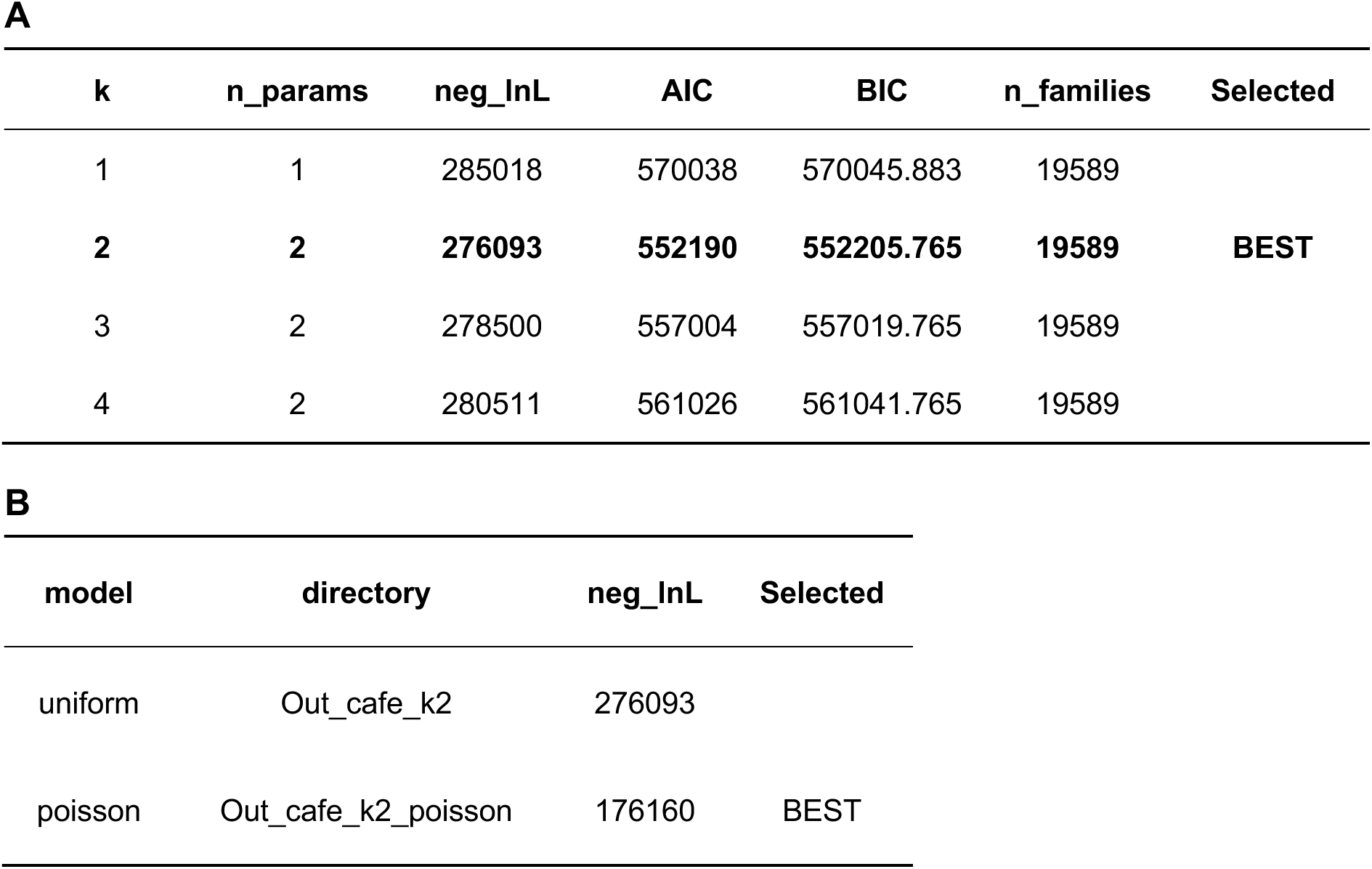
CAFE5 gamma model selection for the 24-species primate dataset. A. Results of CAFE5 cross comparison across increasing numbers of discrete gamma rate categories (k). n_params: number of free parameters estimated; neg_lnL: negative log-likelihood; AIC: Akaike Information Criterion; BIC: Bayesian Information Criterion; n_families: gene families included in the analysis. The best-fit model (lowest AIC) is indicated. k=1 is equivalent to a single global birth-death rate with no among-family rate variation. **B.** Once we have selected, a further CAFE5 run was performed under a Poisson root-frequency distribution and compared against the uniform-prior result already obtained during the k sweep.

Across the phylogeny, gene gain and loss were unevenly distributed (**Figure 2**; **Supplementary Table S2; Supplementary Figure 1**). Considering every family with any net copy-number change (**Supplementary Table 2**, *_HOGs_total columns), most lineages (19 of 24 species) showed more contracting than expanding families, and *Homo sapiens* showed the largest imbalance (236 expansions vs 1,162 contractions). Restricting to statistically significant changes (**Supplementary Table S2**, *_HOGs_significant columns, p ≤ 0.05) reversed this pattern: 21 of 24 species showed more significant expansions than contractions, and 22 of 24 showed a net gain in total genes rather than a net loss. This reversal held at stricter thresholds too (**Supplementary Table S3**): 17 of 24 species at p ≤ 0.01 and 20 of 24 at p ≤ 1×10^−5^ still showed more significant expansions than contractions*. H. sapiens* was a clear exception, showing a significant net contraction at every threshold tested (94 vs 44 families at p ≤ 0.05, down to 9 vs 7 at p ≤ 1×10^−5^; **Supplementary Table S3**), as did *Hylobates moloch* by total genes despite a near-even split in family counts. *Trachypithecus francoisi,* by contrast, showed a consistent excess of expansion by both family count and genes gained across all three thresholds (**Supplementary Table S3**), suggesting genuine lineage-specific diversification rather than a raw-count artefact.

**Figure 1.**
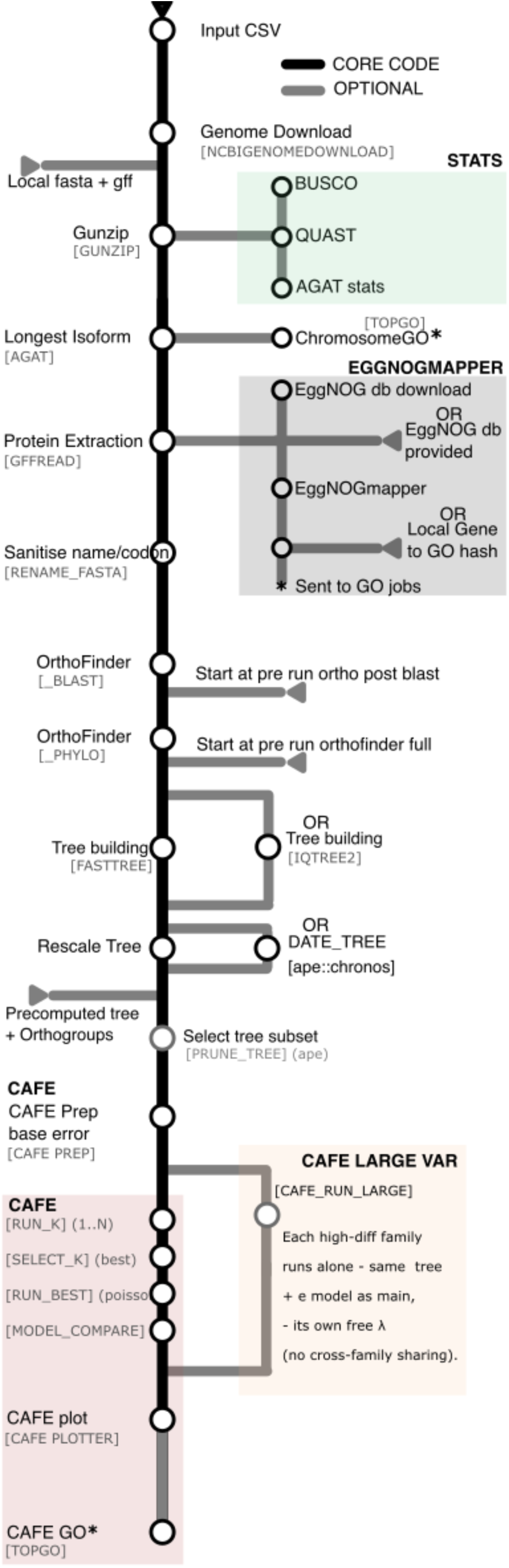
A train map of the main steps in the pipeline. Nodes are the programs/modules that run, and lines show the connections between each program. Grey lines represent optional routes, and grey triangles are optional inputs. The black triangle represents the input samplesheet required to initiate the pipeline. Coloured boxes represent grouped modules related to a specific goal (e.g. CAFE analysis, or the quality control genome statistics [‘STATS’]).

**Figure 2.**
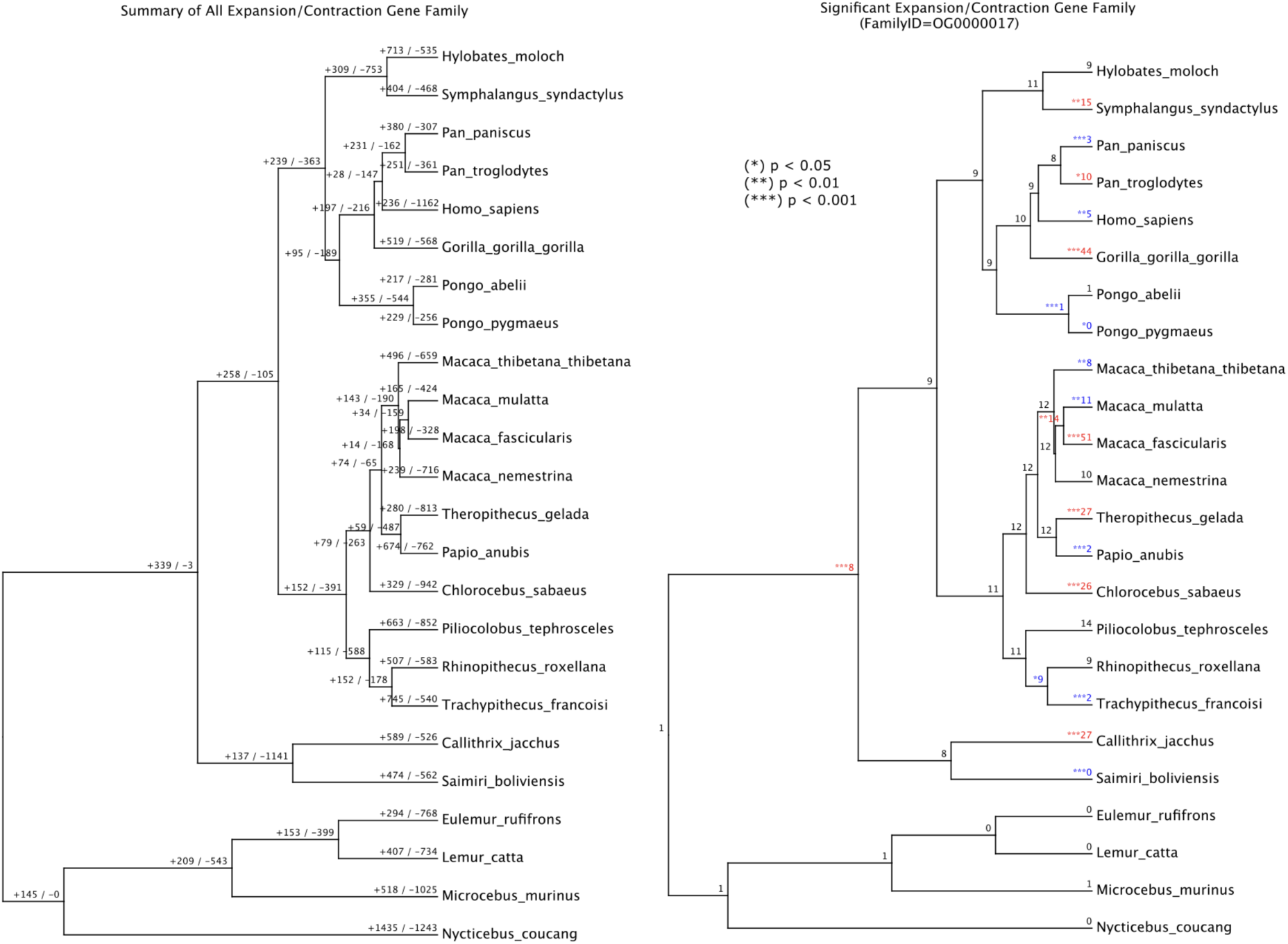
Phylogenetic expansion and contraction trees of primate species. Left: results of CAFE5 showing the number of expanded and contracted gene families at each leaf and node (all families, regardless of significance). Right: a “cafeplotter” tree for orthogroup OG0000017, a DUX (double homeobox) family gene residing in the D4Z4 macrosatellite repeat, showing gene copy number and significance at each node/leaf, for expansion (red) and contraction (blue); (*) p < 0.05, (**) p < 0.01, (***) p < 0.001.

At a family-wide p-value threshold of 0.01, 358 gene families (orthogroups) showed significantly elevated rates of expansion or contraction in at least one species (1,029 significant species-within-orthogroup instances; **Supplementary Table S3**). One of the most dynamic families in the dataset was OG0000017 (significant at 18 species/nodes, p ≤ 0.05), annotated as a member of the DUX (double homeobox) gene family, PRD-class homeodomain transcription factors that reside in the D4Z4 macrosatellite repeat array. Rather than a single lineage-specific event, OG0000017 showed independent, statistically extreme copy-number changes across much of the primate tree, in both directions with strong expansions in *Macaca fascicularis* (+37 copies, p = 1.33×10^−43^), *Gorilla gorilla* (+34, p = 2.16×10^−31^), *Theropithecus gelada* (+15, p = 7.26×10^−15^) and *Callithrix jacchus* (+19, p = 1.35×10^−11^), alongside strong contractions in *Papio anubis* (−10, p = 9.59×10^−13^), *Trachypithecus francoisi* (−7, p = 6.09×10^−8^) and *Saimiri boliviensis* (−8, p = 3.69×10^−7^). This bidirectional, lineage-independent volatility matches the known biology of the DUX4/D4Z4 locus: a macrosatellite repeat known to undergo recurrent, independent structural restructuring across primates via unequal recombination, recently documented directly across a comparable set of primate genome assemblies(40). D4Z4-type tandem repeats are also a recognised source of genome assembly error, so some of this signal may reflect assembly quality differences between species rather than purely biological turnover; nonetheless, recovering a known, independently-verified hotspot of primate copy-number instability from a blind phylogenomic analysis, without prior family curation, is a meaningful demonstration that EXCON’s gene family calls track genuine evolutionary signal. We are not aware of any previously published CAFE-based analysis of this family across a comparable primate dataset.

Gene Ontology enrichment revealed a more structured pattern than a simple gain/loss split (**Figure 3; full enrichment results in Supplementary Table S4**). Terms related to core translation machinery, ribosome, ribosomal subunit, translational initiation, peptide biosynthesis, were broadly enriched among expanding families across most of the phylogeny, rather than contracting families. A narrower, more specific signal was found among contracting families: mitochondrial and oxidative-phosphorylation terms (aerobic and respiratory electron transport chain, ATP synthesis) were significantly enriched, but only in *H. sapiens*, *M. nemestrina* and *T. gelada*, not broadly across the clade. The clearest lineage-specific signal was chemosensory: olfactory receptor activity, odorant binding and detection of chemical stimulus were strongly and almost exclusively enriched among expanding families in *Eulemur rufifrons*. This is consistent with the substantially larger functional olfactory receptor gene repertoire retained in strepsirrhine primates (600–800 genes) compared to haplorhines (200–400 genes), reflecting their greater reliance on olfaction over vision(41).

**Figure 3.**
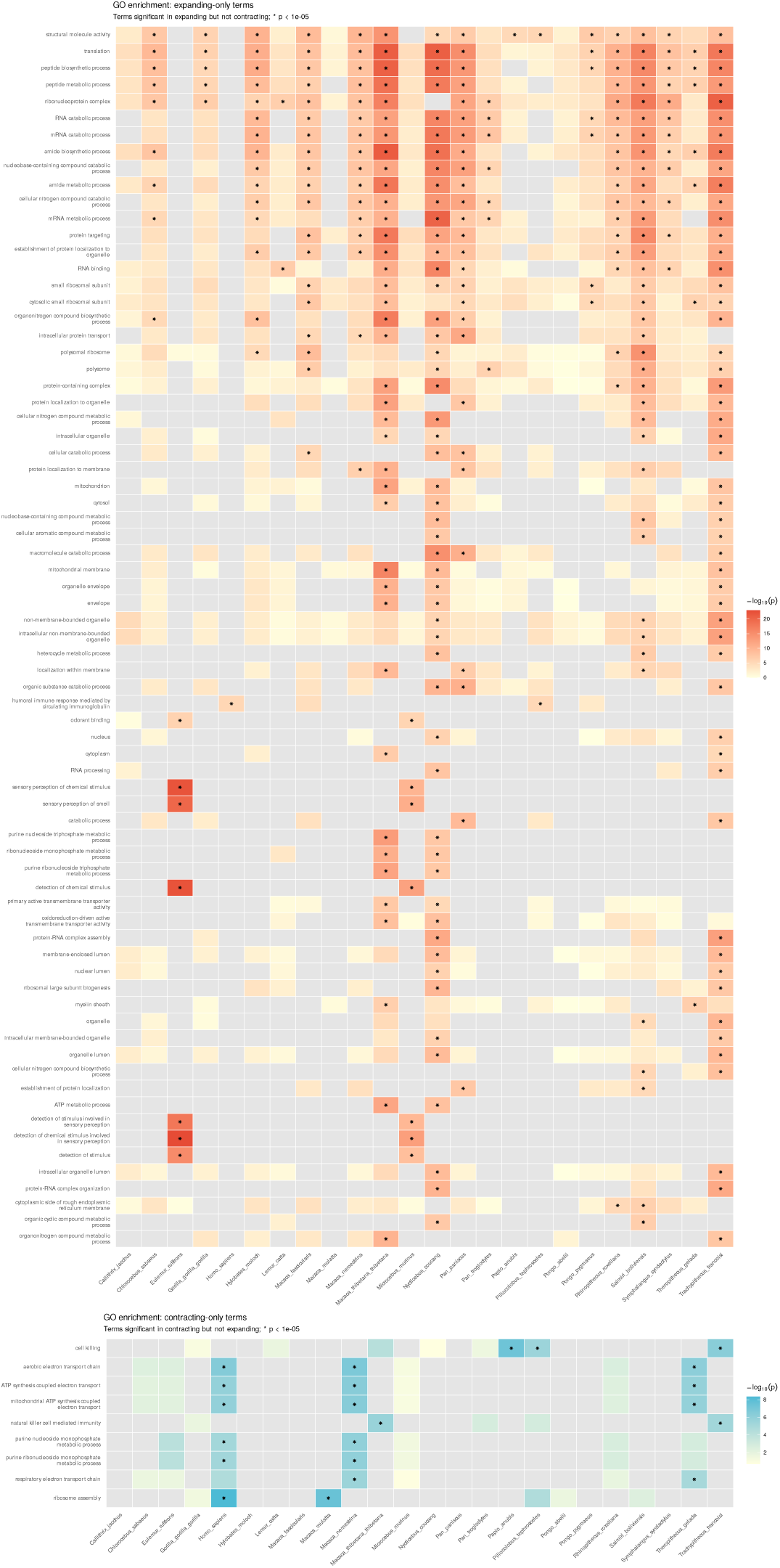
GO term enrichment in expanding and contracting gene families shared across primates. Heatmap showing GO terms significantly enriched (p < 0.00001) in gene families both expanding (top) and contracting (bottom) in at least two primate species. Colour intensity represents the –log_10_(p-value) from CAFE5 gene family evolution GO analysis, with grey indicating no significant enrichment detected in that species. Asterisks (*) denote species where the term met the significance threshold (p < 0.00001). GO terms are ordered by the total number of significant species across both expanded/contracted sets.

### 2Performance and scalability

To ensure the pipeline works on a variety of input data types, we ran the central components of the EXCON pipeline on groups represented in bacteria, insects (just Hymenoptera) and mammals, using a benchmarking dataset, included in the repository (https://github.com/Eco-Flow/excon/tree/main/benchmark). The benchmark runs with a series of commands documented in its README.md, which compare wall time, CPU and memory usage across runs.

With respect to wall time (the actual physical time elapsed between the start and end of the program; **Figure 4**), the pipeline is dominated by OrthoFinder (blast and phylo steps), with CAFE5 taking up the next largest share of time. Occasionally, AGAT can run out of memory and must be resubmitted, which accounts for the additional time in some mammal runs (**Figure 4**: larger green bars; AGAT). Peak memory use reached 50.6GB during OrthoFinder’s phylogenetic inference step (mammal, n=50); AGAT’s longest-isoform selection was the next largest consumer, at up to 15.5GB for large mammalian genomes (**Supplementary Table S5**). All other steps stayed under 6GB.

**Figure 4.**
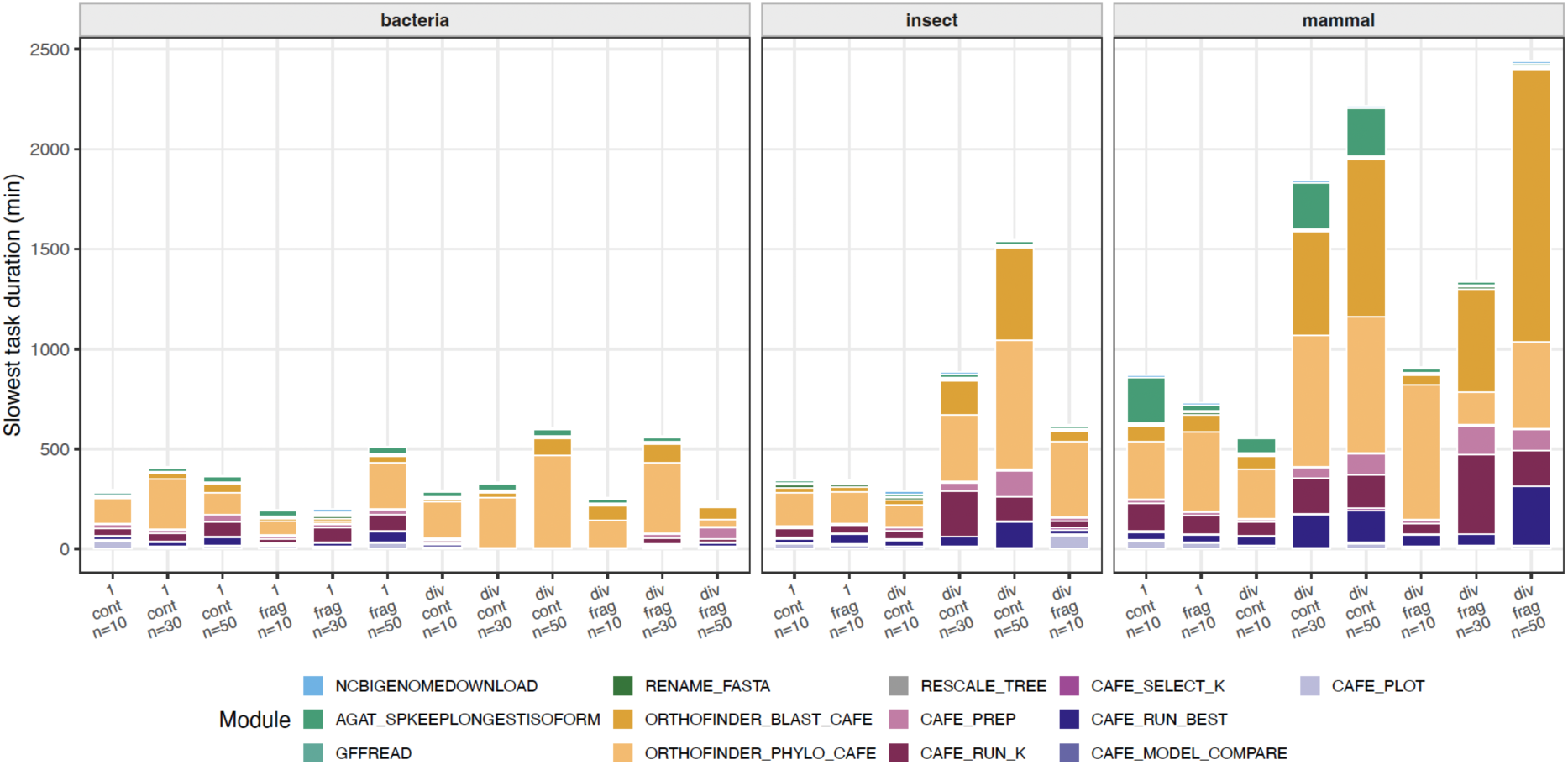
Benchmarking of wall time in different runs. In bacteria, insects (hymenopera) and mammals, we show for each run (bar), the duration of the whole pipeline split into individual modules. Where there is parallelisation, we only show the slowest task duration of one parallel process. Runs are denoted ‘1’ if they came from closer phylogenetic distance (Streptococcus, Apidae, Primates), or ‘div’ (diverse; Alphaproteobacteria, Hymenoptera, Mammals). ‘cont’ is for contiguous genomes, and ‘frag’ is for more fragmented genomes. The number of samples per run is denoted (‘n=’).

Next, we checked if each benchmarked dataset ran, and compared how the runs were affected by the genome size and N50 (a measure of contiguity) (**Figure 5**). These were measured using the stats from the (‘--stats’) QC flag within the EXCON pipeline, which provides average scores for N50 and genome length. In this way, we could plot the passing of successful and failed runs (**Figure 5**). This showed that CAFE will mostly complete across all the datasets we tested, but we did have some failures with the diverse and tiny genome bacterial group “Alphaproteobacteria” (noted as X in **figure 5**). The run times were relatively similar between “close” and “diverse” inputs, suggesting more diverse genomes do not increase the time needed to process the data. Using this information (**Figure 5**), users can estimate the likelihood that their particular samples of interest may pass successfully, as well as the amount of time it may take to process their clade (**Figure 4**).

**Figure 5.**
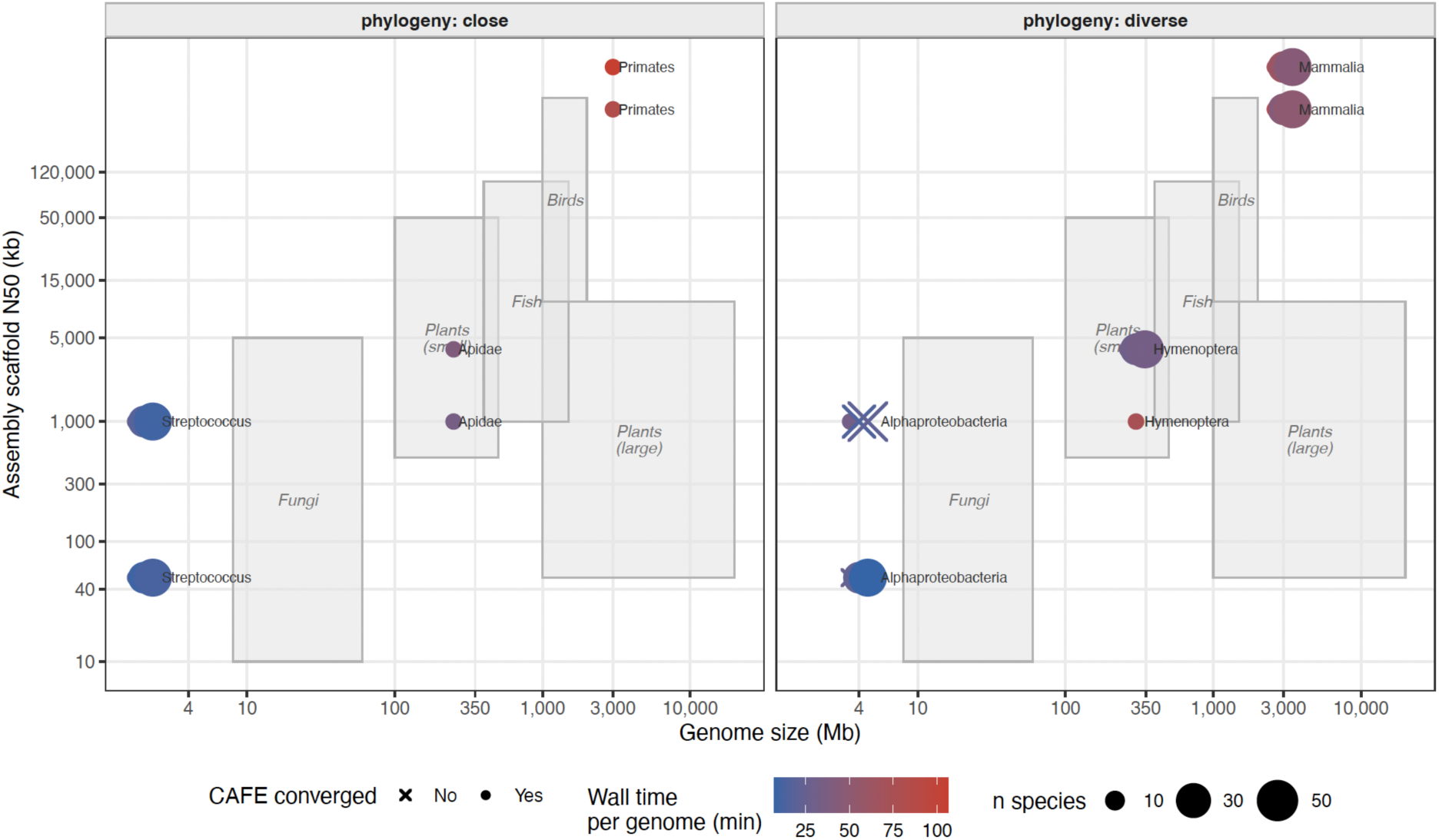
A plot comparing successful and failed EXCON runs. Unsuccessful runs are indicated by an X (CAFE failed to converge). Input datasets contained 10, 30 or 50 genomes (depending on clade availability in RefSeq), with either close or diverse phylogenetic distance (left/right panels respectively). Runs are plotted by representative genome size (Mb, x-axis) and assembly contiguity (N50, y-axis). Point colour shows wall time per genome (minutes); point shape shows whether CAFE5 converged. Grey boxes show typical genome size/N50 ranges for other organism groups, for orientation. Users can locate their own organism relative to the labelled clades to estimate expected run time and the likelihood of CAFE5 convergence.

## Discussion

Identifying gene-family expansions and contractions across species is a core analysis for evolutionary biology. With rapidly expanding genomic resources, there is a need to ensure that such core analyses are conducted to the same high standard in a reproducible manner, and that such methods are accessible to the community, not just those with access to an expert bioinformatician. EXCON addresses a genuine need in comparative genomics for an accessible, reproducible, end-to-end pipeline for gene family evolution analysis. By integrating genome acquisition, protein extraction, orthology inference (OrthoFinder), tree preparation, gene birth-death modelling (CAFE), and functional annotation (eggNOG-mapper) into a single Nextflow workflow, EXCON eliminates the most common sources of error and irreproducibility in such analyses: these include manual file format conversions, version control, ad hoc parameter choices, and undocumented intermediate steps.

Several design choices therefore deserve consideration.

The choice of how to handle sequences containing internal stop codons can affect inferred gene-family expansions and contractions. Removing such sequences (-- internal_stop_action drop) may exclude genuine genes, whereas retaining only the longest ORF (longest_orf) may produce an incomplete sequence representing only a conserved domain, potentially preventing it from clustering with the corresponding gene family. The most appropriate strategy may therefore vary between species, and testing alternative stop-codon handling approaches may be necessary.

The progressive differential filtering strategy is a pragmatic response to the reality that CAFE5’s sensitivity to high-variance gene families means that analysis of unfiltered data frequently fails to converge, particularly on phylogenetically diverse datasets. Our strategy of halving the threshold on each retry minimises the number of families removed while increasing the chance of convergence. Similarly, preserving the removed families for fixed-lambda re-analysis ensures that the most biologically interesting families are those with the most extreme copy number variation and are not simply discarded but are still subject to evolutionary inference. However, we should caution that having separate models for a specific family means that we cannot truly compare these with the other families under a shared model. Ideally in future versions of this pipeline we can address this issue in a more integral way, rather than allowing separate inference of high variance families.

The automatic k selection via AIC is an important addition to the standard CAFE5 workflow. Published analyses frequently use a fixed k without justification, yet the optimal k varies substantially across datasets. EXCON’s parallel k-sweep adds minimal computational overhead (the six runs are run in parallel) and provides a principled basis for model choice.

EXCON currently advises on the use of chromosome-level assembly as a requirement (in the documentation) and optionally reports BUSCO completeness to help users assess assembly quality; however, on slightly poorer contiguous assemblies (e.g. the Hymenoptera and mammalian benchmark data, n=10) our pipeline still successfully runs CAFE. The strategy to filter highly variable orthogroups was also a useful method to “force” CAFE to finish successfully.

However, it should be noted that using poorly contiguous assemblies might inflate spuriously the number of gained and lost genes copies on specific branches of the tree. Nevertheless, poorly contiguous assemblies may spuriously inflate the number of gene copies inferred to have been gained or lost on particular branches of the tree, as has also been reported in CAFE analyses(42). The use of such assemblies should therefore be treated with caution.

Future development of EXCON will focus on: i) additional tooling that can be helpful alongside the given tools (e.g. iqtree3 fossil calibration); ii) KEGG terms; iii) transposable element pre-filtering; and iv) expanded visualisation outputs. All of which are in the roadmap to be tackled next.

EXCON was developed under the BBSRC funded Eco-Flow initiative at University College London, which aims to build a community of researchers producing and sharing reproducible, scalable bioinformatics pipelines for ecology and evolutionary biology. We encourage community contributions, feature requests, and adoption by the broader comparative genomics community (https://github.com/Eco-Flow/).

## Conclusion

To our knowledge, EXCON provides the first fully automated, portable, and reproducible pipeline for end-to-end gene family expansion and contraction analysis. By combining genome acquisition, orthology inference, ultrametric tree construction, CAFE5 model selection, and GO enrichment in a single Nextflow workflow, EXCON makes rigorous analysis accessible to researchers without specialist bioinformatics expertise. Using a progressive convergence recovery strategy, automatic gamma model selection, and error model estimation specifically helps manage this complex analysis. EXCON is freely available and community-driven, and we anticipate that it will become a standard tool for comparative genomics studies across the tree of life.

## Data availability

EXCON is available at https://github.com/Eco-Flow/excon under the MIT licence. The version of this paper is v2.5.0. Test datasets are available within the github repository itself, and are simply lists of Refseq IDs all contained within the repository. For example, the 24-species Primate analysis are available in the repository (https://github.com/Eco-Flow/excon/blob/dev/conf/primate.config), and run with ‘nextflow run main.nf -profile primatè to replicate the results.

## Acknowledgements

We thank the developers of CAFE first and foremost. CAFE is the scientific core of this work; EXCON is best understood as a pipeline built around it and the other major tools it depends on, particularly OrthoFinder for orthology inference, which all require citation if used in the pipeline. We thank members of the Sumner lab (UCL) and Tom Wenseleers lab (KU Leuven) for testing and suggesting improvements to the pipeline, and the wider nf-core community for providing some of the modules we use from nf-core. We thank the UCL Research Computing team for access to the Myriad HPC cluster which permitted extensive testing of the pipeline.

## Contributions

Christopher Wyatt: Conceptualization, Methodology, Software, Formal analysis, Data curation, Writing – original draft, Visualization, Project administration. Alexandra Cerqueira De Araujo: Conceptualization, Writing – review & editing. Fernando Duarte: Code editing. Simon Murray: Code editing. Cintia Oi: Writing – review & editing. Seirian Sumner: Supervision, Funding acquisition. All authors reviewed and approved the final manuscript.

## Conflict of interest

The authors declare no competing interests.

**Supplementary Table S1**. Premature in-frame stop codons resolved during preprocessing, one file per species (24 files). Columns: gene_id, action (longest_orf, the pipeline default for this run), original_length, kept_length (both in amino acids).

**Supplementary Table S2.** Per-species and per-internal-node summary of CAFE5-inferred gene family expansion and contraction. Columns: Total/Expansion/Contraction_HOGs_significant (families with a branch p-value ≤

0.05), Expansion/Contraction_genes_significant (net genes gained/lost among significant families), and Expansion/Contraction_HOGs_total (families with any change, regardless of significance). Internal node numbers correspond to Supplementary Figure S1.

**Supplementary Table S3.** Expansion/contraction summary at stricter significance thresholds (p ≤ 0.01, p ≤ 1×10^−5^), as a two-tab workbook. “Summary” tab: per-species family and gene counts at each threshold. “Raw data” tab: the full per-family, per-node significant changes (Node, HOG, change, pvalue, direction) underlying that summary.

**Supplementary Table S4.** Full GO term enrichment results underlying Figure 3, covering all 24 species (not restricted to the terms shown in the figure). Go_summary_pos.tsv and Go_summary_neg.tsv give per-species enrichment p-values for expanding and contracting families respectively; Go_summary_posneg_merged.tsv combines both; go_enrichment_summary.tsv summarises the number of significant species per GO term and direction.

**Supplementary Table S5.** Peak memory usage per pipeline process across all 26 benchmark runs (Figure 4, Figure 5). Columns: n_runs (number of benchmark runs that process appeared in), mean_peak_ram_gb, max_peak_ram_gb, and run_at_max (which run reached that maximum).

---

**Supplementary Figure S1.**
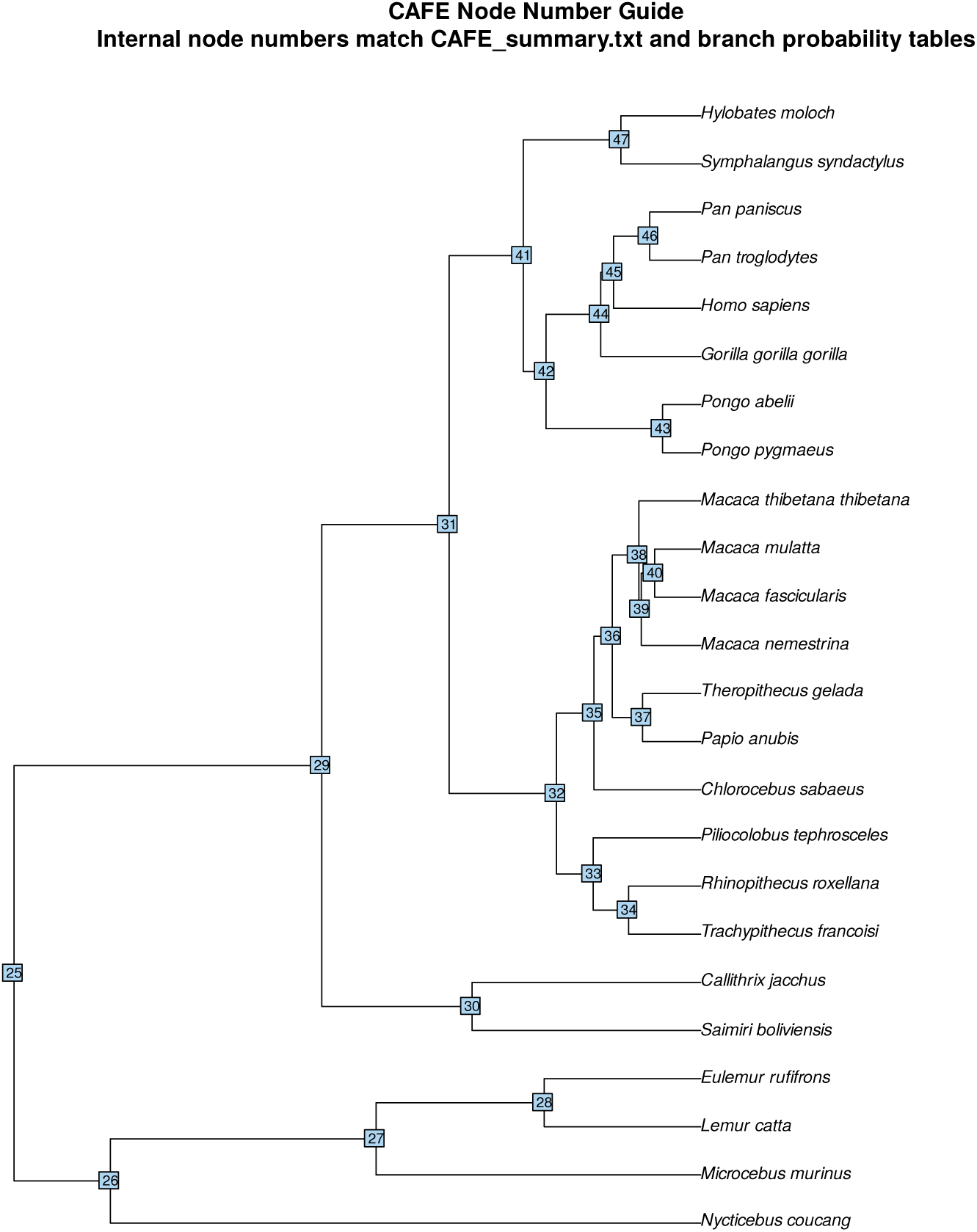
Node numbering guide for the 24-species primate phylogeny. Internal node numbers correspond to the “Species/Node” column in Supplementary Table S2 and to node labels used throughout the CAFE5 output

